# Safety evaluation of soy leghemoglobin protein preparation derived from *Pichia pastoris*, intended for use as a flavor catalyst in plant-based meat

**DOI:** 10.1101/196766

**Authors:** Rachel Z. Fraser, Mithila Shitut, Puja Agrawal, Odete Mendes, Sue Klapholz

**Author notes:** Correspondence to: Rachel Z. Fraser, Impossible Foods Inc., 525 Chesapeake Drive, Redwood City, CA, 94063, USA.

## Abstract

The leghemoglobin protein (LegH) from soy (*Glycine max*) expressed in *Pichia pastoris* (LegH Prep) imparts a meat-like flavor profile onto plant-based food products. The safety of LegH Prep was evaluated through a series of *in vitro* and *in vivo* tests. The genotoxic potential of LegH Prep was assessed using the bacterial reverse mutation assay (Ames test) and the *in vitro* chromosome aberration test. LegH Prep was non-mutagenic and non-clastogenic in each test, respectively. Systemic and female reproductive toxicity were assessed in two separate 28-day dietary studies in Sprague Dawley rats. There were no mortalities associated with the administration of LegH Prep. There were no clinical observations, body weight, ophthalmological, clinical pathology, or histopathological changes attributable to LegH Prep administration. Female reproductive parameters were comparable between rats treated with LegH Prep and concurrent control rats. These studies establish an NOAEL of 750 mg/kg/day LegH, which is over 100 times greater than the 90^th^ percentile estimated daily intake (EDI). Collectively, this work demonstrates that LegH Prep is safe for its intended use in ground beef analogue products at concentrations up to 0.8% LegH.

Abbreviations

LegH: soy leghemoglobin protein
LegH Prep: soy leghemoglobin protein preparation derived from *Pichia pastoris*

## Introduction

Western diets containing meat have a larger negative impact on the environment compared to plant-based diets.^1–4^ However, due to both social and personal reasons, many consumers are resistant to reducing the amount of meat they eat.^2,5,6^ To date, plant-based diets have been limited to small, motivated populations, such as consumers who follow vegetarian or vegan principles.^7–9^ One potential way to catalyze widespread shift to more sustainable, plant-based diets is to create meat directly from plants that satisfies the tastes of meat consumers.^7,8,10^ Achieving that goal would require products that recreate the sensory properties that people crave in meat, including texture, mouthfeel, taste, smell, and cooking experience, based on an understanding of the biochemical origins of meat sensory attributes.

An investigation of the molecular mechanisms underlying the unique flavors and aromas of meat led to the discovery that heme is the critical catalyst of the chemical reactions that transform simple biomolecules into the complex array of odorants and flavor molecules that define the characteristic flavor profile of meat.^11^ Heme is an iron-containing porphyrin ring that exists as a protein co-factor in all branches of life and is essential for most biochemical processes involving molecular oxygen.^12^ Myoglobin and hemoglobin proteins from animal meat tissues have been consumed throughout human history and represent an important source of dietary iron.^13,14^ Plants contain symbiotic and non-symbiotic hemoglobin proteins, both of which share a common ancestor with animal hemoglobins.^15^ Symbiotic plant hemoglobins, also known as leghemoglobins, are present in the root nodules of leguminous plants.^16^ Leghemoglobin controls the oxygen concentration in the area surrounding symbiotic nitrogen-fixing bacteria.^15–17^ Non-symbiotic plant hemoglobins are expressed in the stems, roots, cotyledon, and leaves and are involved in oxygen homeostasis pathways.^18,19^ Due to the presence of non-symbiotic hemoglobins in legumes, cereals, and other plants,^12,15,20^–22 low levels of plant heme proteins are widely consumed in the human diet.^23–25^

Paralleling the catalytic activity of myoglobin in muscle meats, leghemoglobin protein (LegH) from soy (*Glycine max*) imparts meat-like flavors and aromas on to plant-based meat products.^11^ While the primary amino acid sequence of LegH is highly divergent from the sequence of animal hemoglobins and myoglobins, the three dimensional structure is highly similar.^26^ Additionally, the heme co-factor bound to LegH (heme B) is identical to heme found in animal meat, which has a long history of safe use in the human diet.^14^ The iron from LegH has an equivalent bioavailability to iron from bovine hemoglobin when supplemented in a food matrix.^27^ However, because soy root nodules, and thus LegH protein, are not commonly consumed in the human diet, the safety properties of LegH remain not fully understood.

In pursuit of the intended use in ground beef analogue products, the gene encoding a soy LegH protein was introduced into the genome of the yeast *Pichia pastoris*, enabling production of high levels of soy leghemoglobin protein preparation (LegH Prep). The total protein fraction of LegH Prep contains at least 65% LegH. The remaining proteins are from the Pichia host. *Pichia pastoris* is non-toxigenic and non-pathogenic and has been used in the recombinant expression of both GRAS and FDA-approved proteins.^28,29^ Here, we evaluated the safety of LegH Prep with both *in vitro* and *in vivo* models. To evaluate potential genotoxicity and carcinogenicity, a bacterial reverse mutation test (Ames test) and an *in vitro* chromosome aberration test were performed. These *in vitro* models showed that LegH Prep is neither mutagenic nor clastogenic. Systemic toxicity was evaluated by a 28-day feeding study in Sprague Dawley rats. An additional 28-day feeding study was performed to evaluate the female estrous cycle and reproductive health. These *in vivo* models demonstrated no adverse effects attributable to the consumption of LegH Prep at the maximum dose tested, which was more than 100 times greater than the 90^th^ percentile estimated daily intake (EDI) in ground beef analogue products. Collectively, these results demonstrate that LegH Prep is safe for its intended use in ground beef analogue products.

## Materials and Methods

### Test Article Production and Analysis

LegH was recombinantly expressed in *Pichia pastoris* during submerged fed-batch fermentation and isolated using filtration-based recovery with food- or pharmaceutical-grade materials.^30^ The Pichia production strain (MXY0291) is derived from a non-toxigenic and non-pathogenic, well-characterized strain lineage that has a history of safe use in manufacturing proteins for use in food and pharmaceuticals.^28,29,31^ This process is compliant with the Enzyme Technical Association’s guidelines for fermentation-produced microbiologically-derived proteins and follows current Good Manufacturing Practices (cGMP).^32,33^ LegH Prep is a frozen liquid and the entire final preparation was used for all *in vitro* safety tests performed in this study. To aid with homogeneous mixing into the animal diet, LegH Prep was freeze-dried prior to use for all *in vivo* studies.

During each of the *in vivo* studies described below, the LegH concentrations within the neat test substance and animal feed samples were analyzed by high performance liquid chromatography (HPLC) to evaluate test article concentration, stability, and homogeneity. LegH Prep was extracted from the feed by adding 50 mM potassium phosphate pH 7.4, 150 mM sodium chloride to each feed sample followed by 1 hour of end-over-end rotation. HPLC was performed using an Agilent 1100 Series instrument with an ACQUITY xBridge BEH125 SEC 7.8 ×150 mm ID 3.5 μm column (Waters). LegH concentration was determined by integration of the 415 nm absorbance at the LegH retention time.

### Estimated daily intake (EDI)

Ground beef analogue products will be formulated to contain approximately the same amount of heme as beef. This equates to a typical and maximum usage rate of 0.6% and 0.8% LegH. The EDI for LegH within ground beef analogue products was calculated based on 100% capture of the U.S. ground beef market, which is approximately 500 times higher than the market size for all meat and poultry analogue products.^a^ The national mean daily consumption of ground beef for males and females ages 2 and over is 25 grams per day (59 g beef/person/day ^*^ 42% of beef sales are ground beef).^34,35^ Replacement of ground beef with ground beef analogue on a 1-for-1 basis would result in typical (0.6% LegH use rate) and maximum (0.8% LegH use rate) LegH EDIs of 150 and 200 mg/person/day, respectively. In accordance with FDA guidelines, the 90^th^ percentile EDI was calculated as two times the maximum EDI or 400 mg/person/day LegH, which corresponds to 6.67 mg/kg bodyweight/day assuming an average body weight of 60 kg. The 90^th^ percentile was used as a basis for safety testing.

### Bacterial Reverse Mutation Assay (Ames Test)

The Ames test (reverse mutation test)^38^ was performed by Product Safety Labs (PSL) (Dayton, NJ) and was conducted in accordance with U.S. Food and Drug Administration GLP regulations (21 CFR Part 58), and the OECD Principles of GLP ENV/MC/CHEM(98)17.^39^ The assay design was based on OECD Guideline 471^40^ and ICH Guidelines S2A and S2B.^41^ Five bacterial strains were evaluated (*Salmonella typhimurium* (ST) TA98, TA100, TA1535 and TA1537 and *Escherichia coli* (EC) WP2uvrA (Molecular Toxicology, Inc., Boone, NC)) according to the plate incorporation and pre-incubation methods in both the presence and absence of a metabolic activation system (S9 mix). Sterile water served as the negative control, while five mutagens including sodium azide (NaN3), ICR 191 acridine, daunomycin, methyl methanesulfonate (MMS) and 2-aminoanthracene (2-AA) (Molecular Toxicology, Inc., Boone, NC) were used as the positive controls. Water was also used as the solvent for the positive controls except for 2-AA which was prepared in dimethyl sulfoxide (DMSO). The initial test followed the plate incorporation method, in which the following materials were mixed and poured over the surface of a minimal agar plate: 100 μL of the prepared test solutions, negative (vehicle) control, or prepared positive control substance; 500 μL S9 mix or substitution buffer; 100 μL bacteria suspension (ST or EC); and 2000 μL overlay agar maintained at approximately 45 °C.

Plates were prepared in triplicate and uniquely identified. Appropriate sterility control check plates (treated with critical components in the absence of bacteria) were included as a standard procedural check. After pouring, plates were placed on a level surface until the agar gelled, then inverted and incubated at 37 ± 2 °C until growth was adequate for enumeration (approximately 65 ± 3 hours).

The confirmatory test employed the pre-incubation modification of the plate incorporation test. The test or control substances, bacteria suspension, and S9 (or substitution buffer) were incubated under agitation for approximately 30 minutes at approximately 37 ± 2 °C prior to mixing with the overlay agar and pouring onto the minimal agar plates before proceeding as described for the initial test. Following incubation, the revertant colonies were counted manually and/or with the aid of a plate counter (Colony Plate Reader: Model Colony-Doc-It^™^). To be considered valid, the background lawn for vehicle control plates had to appear slightly hazy with abundant microscopic non-revertant bacterial colonies. The mean revertant colony counts for each strain treated with the vehicle had to lie close to or within the expected range taking into account the laboratory historical control range. For each experimental point, the Mutation Factor (MF) was calculated by dividing the mean revertant colony count by the mean revertant colony count for the corresponding concurrent vehicle control group. The results were considered to be positive when the MF was increased at least by a factor of two for strains TA98, TA100 and WP2 uvrA or by at least a factor of three for strains TA1535 and TA1537. In addition, any increases had to be dose-related and/or reproducible, i.e., increases must be obtained at more than one experimental point (at least one strain, more than one dose level, more than one occasion or with different methodologies).

### Chromosome Aberration Assay in Human Peripheral Blood Lymphocytes (HPBL)

The chromosomal aberration assay^42^ was conducted at Eurofins Biopharma (Munich, Germany) in compliance with the German GLP regulations according to 19b Abs. 1 chemikaliengesetz^43^ and the protocol procedures described in the Term Tests for Genetic Toxicity and OECD 473, *In Vitro* Mammalian Chromosome Aberration Test^39,44^ and the European Commission Regulation (EC) No.440/2008 B.10.^45^ The study was conducted using human peripheral blood lymphocytes (HPBL) in both the absence and presence of the chemically-induced rat liver S9 metabolic activation system (Trinova Biochem, Giessen, Germany). Peripheral blood lymphocytes were obtained from healthy non-smoking donors who had no recent history of exposure to genotoxic chemicals and radiation. Peripheral blood lymphocytes were cultured in complete medium (RPMI-1640 containing 15% heat inactivated fetal bovine serum, 0.24 g/mL of phytohemagglutinin and 100 units penicillin and 100 μg/mL streptomycin). The cultures were incubated under standard conditions (37 °C in a humidified atmosphere of 5% CO_2_ in air) for 48 hours. The cells were treated for periods of 4 or 24 hours in the non-activated test system and for a period of 4 hours in the S9-activated test system. All cells were harvested 24 hours after treatment initiation. Cyclophosphamide and ethylmethanesulfonate (Sigma-Aldrich, MO) were evaluated as the concurrent positive controls for treatments with and without S9, respectively.

In addition to the mitotic index determination, the proliferation index of selected samples (negative control and high doses of LegH Prep) was calculated using the BrdU (5-bromo-2’-deoxyurindine) technique. The proliferation index was calculated using Equation 1 (where M1 is the first generation, M2 is the second generation, and M3 is the third generation) based on the number of cell divisions undertaken during the experiment.

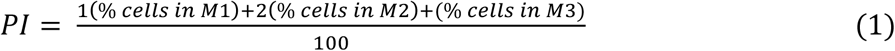

### 14-Day Dietary Palatability and Range Finding Study in Rats

This study^46^ was conducted at PSL (Dayton, NJ) following the OECD 407 Guidelines for Testing of Chemicals^47^ and Food Ingredients and U.S. FDA Toxicological Principles for the Safety Assessment of Food Ingredients IV.C.4.a^48^ and was approved by the Institutional Animal Care and Use Committees (IACUC) of PSL. PSL is AAALAC (Association for Assessment and Accreditation of Laboratory Animal Care) accredited and certified in the appropriate care of all live experimental animals and maintains current staff training ensuring animals were handled humanely during the experimental phase of this study in compliance with the National Research Council’s 2011 Guide for the Care and Use of Laboratory Animals (8^th^ ed.).^49^ CRL Sprague-Dawley CD® IGS rats were purchased from Charles River Laboratories (Kingston, NY) and subsequently quarantined and acclimated to the PSL facilities. Animals were maintained in a temperature- and humidity-controlled room at 19-22 °C and 41-65%, respectively, under a 12 hour light–dark cycle, and fed a standard Envigo Teklad Global 16% Protein Rodent Diet® #2016 (Envigo Laboratories, Inc., Indianapolis, IN). The diet and filtered tap water were supplied *ad libitum*. The animals were group housed and received enrichment activities such as chew sticks throughout the duration of the study. Forty-eight animals were selected for the test (7-8 weeks of age at dosing; weighing 230-264g (males) and 158-181g (females)) and distributed into four groups with 6 males and 6 females each (1 control group per sex and 3 dietary levels per sex). The freeze-dried LegH Prep was administered in the diet. The animals were observed daily for viability, signs of gross toxicity, and behavioral changes at least once daily during the study, and weekly for a battery of detailed observations. Body weights were recorded two times during the acclimation period (including prior to dosing on study day 1) and on study days 3, 7, 10, and 14. Individual food consumption was also recorded to coincide with body weight measurements. Food efficiency was calculated by dividing the mean daily body weight gain by the mean daily food consumption. The animals were fasted overnight prior to blood collection. Samples were collected from all animals for hematology evaluation via the inferior vena cava under isoflurane anesthesia during the necropsy procedure. Approximately 500 µL of blood were collected in a pre-calibrated tube containing K_2_EDTA. All clinical pathology samples were sent to DuPont Haskell Global Centers for Health and Environmental Sciences (Newark, DE) for analysis. Gross necropsy was performed on study day 15 and the animals were evaluated for any macroscopic changes. Histological examination was performed on the liver, spleen, and bone marrow of the animals from the vehicle control and high dose (groups 1, and 4, respectively). Slide preparation was performed by Histo-Scientific Research Laboratories (HSRL) (Mount Jackson, VA) and histopathological assessment was performed by a Board Certified Veterinary Pathologist at PSL.

### 28-Day Dietary Feeding Study in Rats

This 28-day feeding study^50^ was conducted at PSL (Dayton, NJ) in accordance with GLP and follows OECD Guidelines for Testing of Chemicals, Section 4 Health Effects (Part 408): Repeated Dose 90-day Oral Toxicity Study in Rodents^39,51^ and the U.S. FDA Toxicological Principles for the Safety Assessment of Food Ingredients, IV.C.4.a.^48^ and was approved by the IACUC of PSL. Adult CRL Sprague-Dawley CD® IGS rats were purchased from Charles River Laboratories (Kingston, NY) and subsequently quarantined and acclimated to the PSL facilities as described above. Eighty rats were selected for testing, using acceptance criteria described above, and distributed into four groups with 10 males and 10 females per group (1 control group per sex and 3 dietary dose levels per sex). The freeze-dried LegH Prep was administered in the diet.

Prior to study initiation and again on study day 23, the eyes of all rats were examined by focal illumination and indirect ophthalmoscopy. Clinical observations, food consumption, body weight, and food efficiency were evaluated as described above. On study day 22, samples were collected from all animals for hematology, serum chemistry and urinalysis evaluation. Blood samples for hematology and serum chemistry were collected via sublingual bleeding under isoflurane anesthesia. Approximately 500 μL of blood were collected in a pre-calibrated tube containing K_2_EDTA for hematology assessments. The whole blood samples were stored under refrigeration and shipped on cold packs. Approximately 1000 μL of blood were collected into a tube containing no preservative for serum chemistry assessments. At terminal sacrifice, all animals were euthanized by exsanguination from the abdominal aorta under isoflurane anesthesia and blood was collected for evaluation of coagulation parameters. All clinical pathology samples were sent to DuPont Haskell Global Centers for Health and Environmental Sciences (Newark, DE) for analysis. All animals in the study were subjected to a full necropsy, which included examination of the external surface of the body, all orifices, and the thoracic, abdominal and cranial cavities and their contents. Tissues/organs representing systems were collected and preserved in 10% Neutral Buffered Formalin with the exception of the eyes, testes and epididymides, which were preserved in Davidson’s fixative before transfer to ethanol. A subset of tissues/organs were weighed wet as soon as possible after dissection to avoid drying including: adrenal glands, kidneys, spleen, brain, liver, thymus, testes, epididymides, ovaries with oviducts, uterus, and heart. The fixed tissues were trimmed, processed, embedded in paraffin, sectioned with a microtome, placed on glass microscope slides, stained with hematoxylin and eosin, and examined by light microscopy. Histological examination was performed on the preserved organs and tissues of the animals from the vehicle control and high dose groups (Groups 1, and 4, respectively), with the exception of the female reproductive organs, which were evaluated for all dose levels. Slide preparation and histopathological assessment was performed by a Board Certified Veterinary Pathologist at Histo-Scientific Research Laboratories (HSRL) (Mount Jackson, VA). A pathology peer review was performed by a Board Certified Veterinary Pathologist at Regan Path/Tox Services (Ashland, OH) for all female reproductive tissues.

### 28-Day Investigative Study with a 14-Day Pre-Dosing Estrous Cycle Determination

A study was conducted at PSL (Dayton, NJ)^52^ and the protocol was approved by the IACUC of PSL. Adult CRL Sprague-Dawley CD® IGS female rats were purchased from Charles River Laboratories and subsequently quarantined and acclimated at the PSL facilities as described above. Sixty rats were selected for testing using the acceptance criteria described above and animals were distributed into four groups with 15 females per group (1 control group per sex and 3 dietary dose levels per sex). Freeze dried LegH Prep was administered in the diet. The estrus cycle was evaluated daily by vaginal cytology for a period of 14 days prior to administration of the test substance and for the last two weeks of the 28-day period of test substance administration. Estrous cycle stage was not evaluated for the first two weeks of the dosing period to avoid over-manipulating the animals. For each 14-day period, average estrus cycle length was calculated for each animal and subsequently each group. Clinical observations, food consumption, body weight, and food efficiency were evaluated as described above. All animals were subjected to a full necropsy, which included examination of the external surface of the body, all orifices, and the thoracic, abdominal and cranial cavities and their contents. Female reproductive organs were collected and preserved in 10% Neutral Buffered Formalin. The ovaries with oviducts and uterus were weighed wet as soon as possible after dissection to avoid drying. Reproductive tissues were fixed and examined as described above. Estrous cycle stage was recorded for all animals. Histological examination was performed on the preserved organs and tissues for group 1 and group 4 animals. Slide preparation was performed by Histoserv Inc. (Germantown, MD) and histopathological assessment was performed by a Board Certified Veterinary Pathologist at Regan Path/Tox Services (Ashland, OH).

### Statistical Analyses

Mean and standard deviations were calculated for all quantitative data. For the Chromosome Aberration Study^42^, the Fisher’s exact test was used to compare the induction of chromosome aberrations in treated cultures and solvent control. Significance was judged at a probability value of *p*<0.05. Male and female rats were evaluated separately. Body weights, food consumption, urine volume, hematology, blood chemistry, and absolute and relative organ weights, averages and standard deviations were calculated, and analyzed by Bartlett’s test for homogeneity of variances and normality.^53^ Where Bartlett’s test indicated homogeneous variances, treated and control groups were compared using a one-way analysis of variance (ANOVA). When ANOVA was significant, a comparison of the treated groups to control by Dunnett’s test for multiple comparisons was performed.^54,55^ Where variances were considered significantly different by Bartlett’s test, groups were compared using a non-parametric method (Kruskal-Wallis non-parametric analysis of variance).^56^ When non-parametric analysis of variance was significant, comparison of treated groups to control was performed using Dunn’s test.^57^ Statistical analysis was performed on all quantitative data for in-life and organ weight parameters using Provantis™ Version 8, Tables and Statistics, Instem LSS, Staffordshire UK. Clinical pathology was preliminarily tested via Levene’s test^58^ for homogeneity and via Shapiro-Wilk test^59^ for normalcy followed by ANOVA followed with Dunnett’s test.^54,55^

## Results

### Bacterial Reverse Mutation Assay (Ames Test)

The objective of this test was to determine the mutagenic potential of LegH Prep using histidine-requiring strains of *S. typhimurium* (TA98, TA100, TA1535 and TA1537) and a tryptophan-requiring strain of *E. coli* (WP2 uvrA). LegH Prep was evaluated with and without an exogenous metabolic activation (S9 mix) at levels of 23.384, 74, 233.84, 740, 2338.4, 7400, 23,384 and 74,000 μg/plate, which corresponded to 1.58, 5.0, 15.8, 50, 158, 500, 1580, and 5000 μg/plate of the characterizing component, LegH, with the high level being the standard limit for this test.

The mean revertant colony counts for each strain treated with the vehicle were close to or within the expected range, considering the laboratory historical control range and/or published values.^60,61^ The positive control substances caused the expected substantial increases in revertant colony counts in both the absence and presence of S9 in each phase of the test, confirming the sensitivity of the test and the activity of the S9 mix (Table 1). No signs of precipitation or contamination were noted throughout the study. Therefore, each phase of the test is considered valid.

**Table 1:**
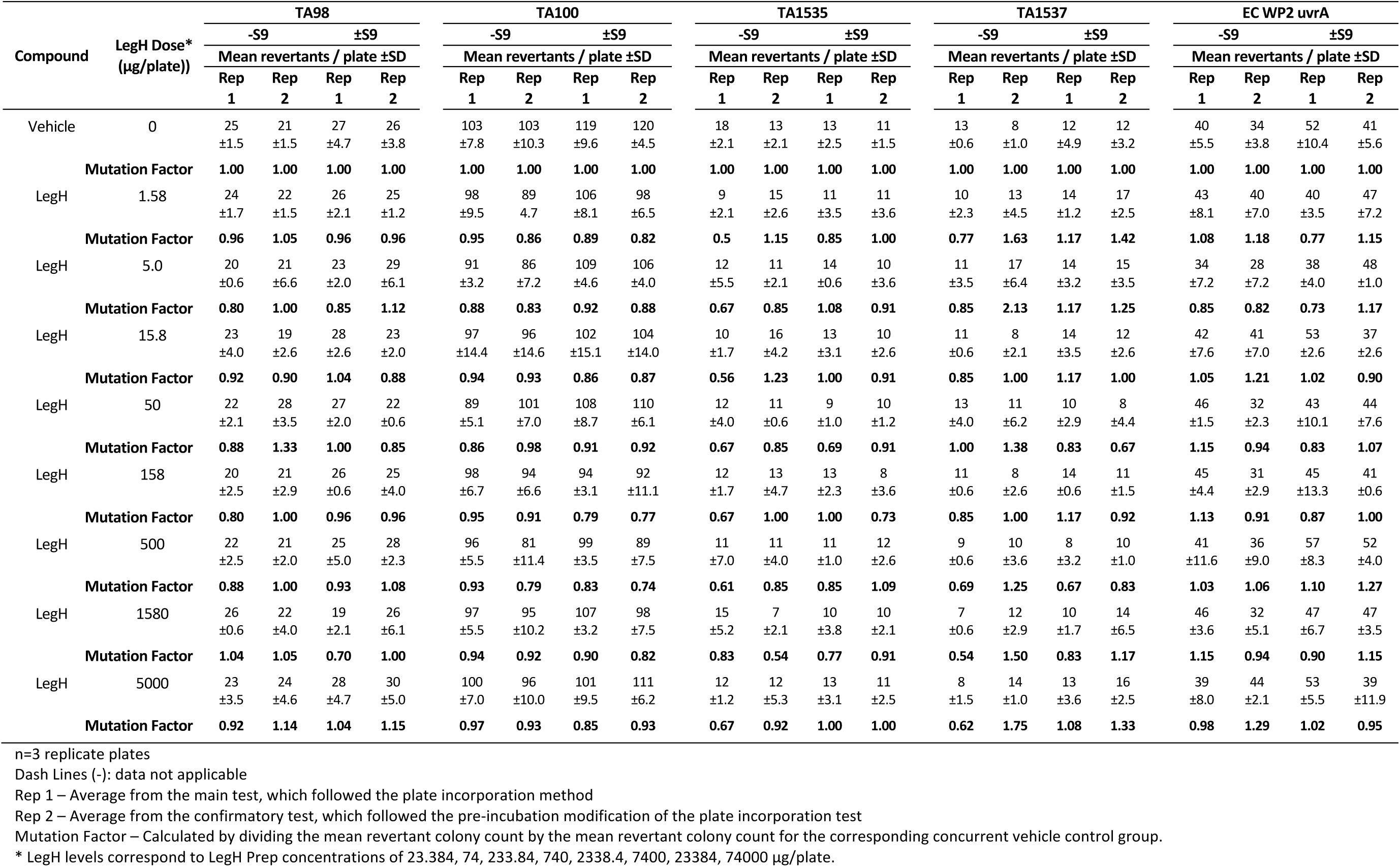
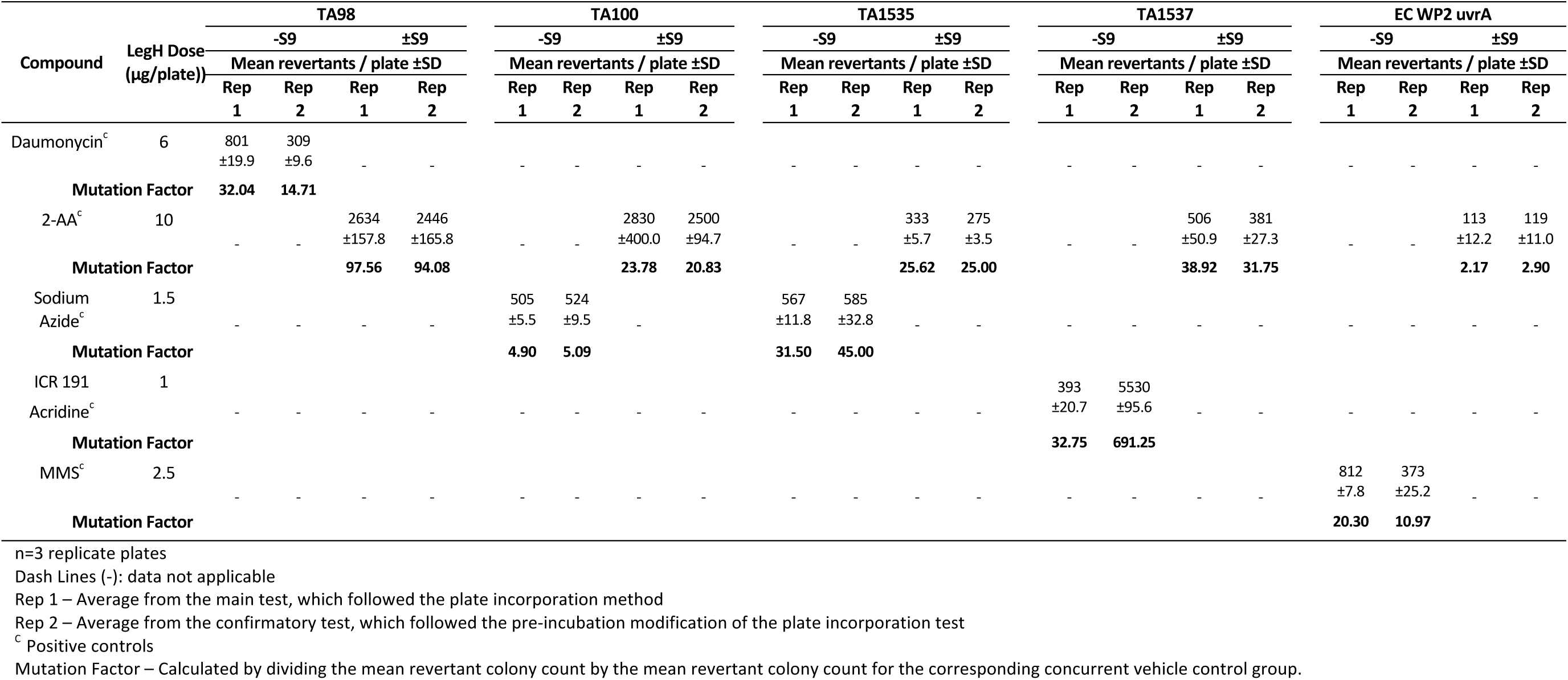
Number of Revertant Colonies and Mutation Factors Without/With Metabolic Activation (S9) - Ames Test.

LegH Prep did not cause a positive increase in the mean number of revertant colonies per plate with strains TA1535, TA1537, TA98, TA100 or WP2 uvrA in either the absence or presence of S9 when using either the plate incorporation or the pre-incubation method (Table 1). Therefore, LegH Prep was non-mutagenic in the bacterial reverse mutation assay.

### Chromosome Aberration Assay in Human Peripheral Blood Lymphocytes (HPBL)

The objective of this *in vitro* assay was to evaluate the ability of LegH Prep to induce structural or numerical (polyploid or endoreduplicated) chromosome aberrations in HPBL. HPBL cells were exposed to LegH Prep for 4 hours in the presence or absence of S9 (Experiment 1) or for 24 hours in the absence of S9 (Experiment 2). In each experiment, untreated and positive controls values were within the historical control range indicating that the subject assay met the criteria for a valid test.

In accordance with OECD guidelines, Experiment 1 used the recommended concentrations of 100-5000 μg/mL LegH, which corresponded to 148-74,000 µg/mL LegH Prep. Mitotic index was evaluated first since decreased mitotic index can inhibit the ability to evaluate chromosome aberrations. Although the mitotic index decreased to below 70% of the negative control at high concentrations of LegH Prep without S9 metabolic activation, no such decrease in the mitotic index was observed in the presence of S9 metabolic activation (Table 2). No significant difference in the proliferation index was observed in either condition (Table 3). Evaluation of chromosomal aberration tests using the mitotic index can be unreliable.^62^ However, in all cases, the mitotic index remained above the 45% of control threshold that is recommended for evaluation of structural and numerical chromosomal aberrations and no test article precipitation was observed. In Experiment 1, no significant increase in cells with structural or numerical chromosome aberrations was observed up to 5000 μg/mL LegH, which was the maximum dose tested, both with and without S9 (Table 2).

**Table 2:** Human Peripheral Blood Lymphocytes Treated with LegH Prep – Chromosome Aberration Assay.

| Treatment<br>(w/v) | S9<br>Activation | Treatment<br>Time | Mean<br>Mitotic Index | Cells<br>Scored | Cells With Aberrations |  |
| --- | --- | --- | --- | --- | --- | --- |
|  |  |  |  |  | Including Gaps | Excluding Gaps |
| Experiment 1 (-S9) |  |  |  |  |  |  |
| Control | -S9 | 4 | 40 | 300 | 17 | 10 |
| LegH 500 (µg/mL) | -S9 | 4 | 35 | 300 | 18 | 12 |
| LegH 1000 (µg/mL) | -S9 | 4 | 28 | 300 | 22 | 13 |
| LegH 2500 (µg/mL) | -S9 | 4 | 23 | 300 | 18 | 12 |
| LegH 5000 (µg/mL) | -S9 | 4 | 22 | 300 | 20 | 5 |
| EMS 900 (µg/mL) | -S9 | 4 | 25 | 200 | 37 | 32 |
| Experiment 1 (+S9) |  |  |  |  |  |  |
| Control | +S9 | 4 | 38 | 300 | 23 | 11 |
| LegH 1000 (µg/mL) | +S9 | 4 | 42 | 300 | 22 | 13 |
| LegH 2500 (µg/mL) | +S9 | 4 | 32 | 300 | 15 | 6 |
| LegH 5000 (µg/mL) | +S9 | 4 | 40 | 300 | 15 | 8 |
| CPA 7.5 (µg/m)L | +S9 | 4 | 34 | 200 | 37 | 31 |
| Experiment 2 (-S9) |  |  |  |  |  |  |
| Control | -S9 | 24 | 60 | 300 | 13 | 7 |
| LegH 100 (µg/mL) | -S9 | 24 | 52 | 300 | 18 | 8 |
| LegH 200 (µg/mL) | -S9 | 24 | 57 | 300 | 20 | 10 |
| LegH 500 (µg/mL) | -S9 | 24 | 42/P | 300 | 10 | 6 |
| LegH 1000 (µg/mL) | -S9 | 24 | 32/P | 300 | 10 | 6 |
| LegH 2000 (µg/mL) | -S9 | 24 | 16/P | ND | ND | ND |
| LegH 3000 (µg/mL) | -S9 | 24 | 8/P | ND | ND | ND |
| LegH 4000 (µg/mL) | -S9 | 24 | 23/P | ND | ND | ND |
| LegH 5000 (µg/mL) | -S9 | 24 | 25/P | ND | ND | ND |
| EMS 400 (µg/mL) | -S9 | 24 | 29 | 75 | 43 | 41 |

ND – Not determined
P - Precipitate observed
<sup>S1</sup> – p <0.05
<sup>S2</sup> – p <0.01
<sup>S3</sup> – p <0.001

**Table 3:** Human Peripheral Blood Lymphocytes Treated with LegH Prep – Proliferation Index.

| Treatment<br>(LegH w/v) | S9<br>Activation | Treatment<br>time (h) | Proliferation<br>Index | Cells in Mitosis Number |  |  |
| --- | --- | --- | --- | --- | --- | --- |
|  |  |  |  | 1 | 2 | 3 |
| Experiment 1 (-S9) |  |  |  |  |  |  |
| Control | -S9 | 4 | 1.16 | 84 | 16 | 0 |
| 5000 (µg/mL) | -S9 | 4 | 1.09 | 91 | 9 | 0 |
| Experiment 1 (+S9) |  |  |  |  |  |  |
| Control | +S9 | 4 | 1.12 | 88 | 12 | 0 |
| 5000 (µg/mL) | +S9 | 4 | 1.07 | 93 | 7 | 0 |
| Experiment 2 (-S9) |  |  |  |  |  |  |
| Control | -S9 | 24 | 1.56 | 44 | 56 | 0 |
| 500 (µg/mL) | -S9 | 24 | 1.23 | 77 | 23 | 0 |
| 1000 (µg/mL) | -S9 | 24 | 1.12 | 88 | 12 | 0 |

<sup>S1</sup> – p <0.05<sup>S2</sup> – p <0.01<sup>S3</sup> – p <0.001

The increased incubation time of Experiment 2 resulted in precipitation of the test article at concentrations of greater than or equal to 500 μg/mL LegH. Additionally, the mitotic index values relative to the control decreased below the 45% percent threshold at concentrations greater than 1000 μg/mL LegH. Therefore, only concentrations up to 1000 μg/mL LegH were evaluated for chromosome aberrations. The proliferation index values for 500 and 1000 micrograms/plate were 1.23 (79% relative to control) and 1.12 (72% relative to control), respectively (Table 3). This decrease was not a consequence of chromosome aberrations. In Experiment 2, no significant increase in cells with structural or numerical chromosome aberrations was observed up to 1000 μg/mL LegH, which was the maximum dose evaluated, without metabolic activation (Table 2).

These results indicate that LegH Prep does not induce structural or numerical chromosome aberrations in either the non-activated or the S9-activated test system. Therefore, LegH Prep is considered non-clastogenic in the *in vitro* mammalian chromosome aberration test using HPBL.

### Animal Feed Analytical Chemistry

In each of the *in vivo* studies described below, the dietary preparations were analyzed using HPLC to evaluate test article (freeze-dried LegH Prep) homogeneity, stability, and concentration verification. In each case, the analyte fell within acceptable parameters: < 10% RSD of LegH concentration between samples of the feed collected from top, middle, and bottom of the mixer, indicating homogeneous distribution; < 10% change in LegH concentration during diet presentation, indicating stability; and within 10% of the target concentration of LegH, indicating accurate dosing.

### 14-Day Dietary Palatability and Range Finding Study in Rats

A 14-day toxicology and palatability feeding study in rats^46^ was performed to assess the feasibility of oral administration of freeze-dried LegH Prep in the diet and to establish the dose range for the subsequent 28-day study. The test article was administered in doses of 0, 3156, 6312, and 12,612 ppm (Groups 1-4, respectively) corresponding to active ingredient LegH concentrations of 0, 125, 250 and 500 mg/kg/day. The calculated nominal dietary intake levels were 134, 269, and 531 mg/kg/day for Group 2-4 male rats and 148, 296, and 592 mg/kg/day for Group 2-4 female rats. The animals are considered to have received acceptable dose levels.

#### Mortality, Clinical Signs, Body Weight/Food Consumption

There were no mortalities during the course of the 14-day study. There were no clinical observations attributed to administration of LegH Prep. In-life clinical signs included reported discoloration of the urine in 6/6 Group 4 males and 5/6 Group 4 females on day 5. Due to its single day occurrence, this observation is likely due to re-hydration of the test article by animal urine, which would result in the formation of a red/brown color. Without a correlation to clinical hematology or any other parameter, these findings are interpreted to be of no toxicological significance. There were no changes in mean food consumption, mean food efficiency, mean body weight, and mean daily body weight gain attributable to the administration of LegH Prep.

#### Pathology

There were no changes in hematology values attributable to the administration of LegH Prep. There were no LegH Prep-related macroscopic or microscopic findings. Mean absolute and relative organ-to-body weights for Group 2-4 were comparable to control group 1 throughout the study. These results suggest that LegH Prep would be well tolerated in a study of longer duration.

### 28-Day Dietary Feeding Study in Rats

A 28-day dietary feeding study in rats was performed to evaluate the potential subchronic toxicity of LegH Prep following continuous exposure of the test substance in the diet. A no observed adverse effect level (NOAEL) was sought for each sex. Due to the successful palatability of 500 mg/kg/day LegH in the previous 14-day feeding study, the maximum dose was increased to 750 mg/kg/day LegH. Administered doses of 0, 512, 1024 and 1536 mg/kg/day of freeze-dried LegH Prep corresponded to 0, 250, 500 and 750 mg/kg/day of LegH, respectively. The slight difference between in correlation between LegH Prep dose levels and LegH concentrations compared to the previous 14-day study are due to the utilization of a different lot of freeze-dried LegH Prep test article. The mean overall daily intake of the test substance in Group 2-4 male rats was 234, 466, and 702 mg/kg/day LegH. The mean overall daily intake in Group 2-4 female rats was 243, 480, and 718 mg/kg/day LegH. The animals are considered to have received acceptable dose levels.

#### Mortality, clinical signs, body weight/food consumption

No mortalities were observed during this study. There were no clinical observations attributable to the administration of LegH Prep. There were no body weight, body weight gain, food consumption, or food efficiency findings considered attributable to LegH Prep administration (Tables 4-6). A statistically significant decrease (p < 0.01) in mean daily body weight gain was observed in group 2 females on days 14-21 (Table 4). This decrease was transient and was interpreted to have no toxicological relevance. Statistically significant increases (p < 0.05-0.01) were observed for mean daily food consumption in Group 3 males on days 7-14 and in Group 4 males on days 7-10, that were transient and without significant impact on body weight, and were interpreted to be non-toxicologically relevant (Table 5). Mean food efficiency for the treated female rats in Group 2-4 was generally comparable to the control Group 1 values throughout the study, with the exception of statistically significant increases (p < 0.01) in Group 2 on Days 14-21 that were transient and without significant impact on body weight and were interpreted to be non-toxicologically relevant. These small but significant changes were all considered to be non-toxicologically relevant and non-test article dependent.

**Table 4:** Summary of Mean Body Weights and Mean Body Weight Gain – 28-Day Dietary Study.

| Mean Body weight (mg/kg/day) |  |  |  |  |  |  |  |  | Mean Body Weight Gain (mg/kg/day) |  |  |  |  |  |  |  |  |
| --- | --- | --- | --- | --- | --- | --- | --- | --- | --- | --- | --- | --- | --- | --- | --- | --- | --- |
| LegH Dose Levels | 0 (mg/kg/day) |  | 250 (mg/kg/day) |  | 500 (mg/kg/day) |  | 750 (mg/kg/day) |  | LegH Dose Levels | 0 (mg/kg/day) |  | 250 (mg/kg/day) |  | 500 (mg/kg/day) |  | 750 (mg/kg/day) |  |
| Study Day | M | F | M | F | M | F | M | F | Study Days | M | F | M | F | M | F | M | F |
| **0** | 236.4<br>±6.1 | 174.1<br>12.3± | 236.4<br>±6.1 | 174.4<br>±12.6 | 236.7<br>±7.0 | 175.6<br>±11.8 | 236.3<br>±6.7 | 174.3<br>±11.9 | **0-7** | 7.33<br>±1.35 | 3.46<br>±1.03 | 7.60<br>±0.94 | 3.80<br>±0.96 | 7.74<br>±1.40 | 4.06<br>±0.72 | 8.07<br>±1.16 | 3.57<br>±0.61 |
| **7** | 287.7<br>±14.0 | 198.3<br>±14.8 | 289.6<br>±11.1 | 201.0<br>±16.5 | 290.9<br>±14.3 | 240.0<br>±13.3 | 292.8<br>±12.2 | 199.3<br>±10.5 | **7-14** | 6.37<br>±0.77 | 2.93<br>±2.09 | 6.77<br>±1.30 | 2.50<br>±0.75 | 7.24<br>±1.49 | 2.81<br>±0.66 | 6.67<br>±1.13 | 3.14<br>±0.81 |
| **14** | 332.3<br>±16.5 | 218.8<br>±21.9 | 337.0<br>±18.4 | 218.5<br>±19.6 | 341.6<br>±24.2 | 223.7<br>±14.6 | 339.5<br>±18.6 | 221.3<br>±14.3 | **14-21** | 5.84<br>±1.15 | 2.91<br>±1.15 | 5.66<br>±0.91 | 1.51<sup>S2</sup><br>±0.82 | 6.13<br>±1.12 | 2.16<br>±1.03 | 5.77<br>±0.86 | 2.39<br>±0.96 |
| **21** | 373.2<br>±22.7 | 239.2<br>±24.0 | 376.6<br>±21.4 | 229.1<br>±19.4 | 384.5<br>±31.1 | 238.8<br>±19.4 | 379.9<br>±22.7 | 238.0<br>±13.1 | **21-28** | 3.07<br>±2.06 | 1.51<br>±1.03 | 3.19<br>±0.75 | 2.13<br>±0.85 | 3.67<br>±1.10 | 2.06<br>±0.63 | 3.66<br>±0.47 | 1.53<br>±0.79 |
| **28** | 394.7<br>±28.8 | 249.8<br>±24.0 | 398.9<br>±26.4 | 244.0<br>±23.3 | 410.2<br>±37.2 | 253.2<br>±17.7 | 405.5<br>±24.4 | 248.7<br>±12.4 | **0-28** | 5.65<br>±0.84 | 2.70<br>±0.60 | 5.80<br>±0.83 | 2.49<br>±0.53 | 6.20<br>±1.14 | 2.77<br>±0.30 | 6.04<br>±0.70 | 2.66<br>±0.34 |

n=10 animals/sex/group
M = male
F = female
<sup>S1</sup> – p <0.05
<sup>S2</sup> – p <0.01
<sup>S3</sup> – p <0.001

**Table 5:** Summary of Mean Daily Food Consumption – 28-Day Dietary Study.

| Mean Daily Food Consumption (mg/kg/day) |  |  |  |  |  |  |  |  |
| --- | --- | --- | --- | --- | --- | --- | --- | --- |
| LegH Dose Levels | 0 (mg/kg/day) |  | 250 (mg/kg/day) |  | 500 (mg/kg/day) |  | 750 (mg/kg/day) |  |
| Study Day | M | F | M | F | M | F | M | F |
| **0-3** | 18.73<br>±3.35 | 13.43<br>±2.05 | 18.73<br>±1.98 | 12.93<br>±2.21 | 18.80<br>±2.68 | 13.73<br>±1.74 | 19.03<br>±2.99 | 13.70<br>±2.56 |
| **3-7** | 28.03<br>±1.08 | 21.18<br>±1.24 | 28.60<br>±0.52 | 21.23<br>±1.13 | 28.23<br>±2.08 | 21.05<br>±1.31 | 28.63<br>±1.07 | 20.18<br>±0.77 |
| **0-7** | 24.04<br>±1.67 | 17.86<br>±0.98 | 24.37<br>±0.65 | 17.67<br>±1.06 | 24.19<br>±0.94 | 17.91<br>±1.02 | 24.51<br>±1.56 | 17.40<br>±0.82 |
| **7-10** | 26.30<br>±1.31 | 19.33<br>±2.23 | 27.10<br>±0.81 | 18.43<br>±0.54 | 27.80<sup>s2</sup><br>±1.97 | 19.30<br>±2.26 | 27.90<sup>s1</sup><br>±0.78 | 18.90<br>±0.96 |
| **10-14** | 26.55<br>±1.17 | 19.55<br>±1.59 | 27.25<br>±0.91 | 20.45<br>±2.02 | 27.88<br>±2.25 | 19.45<br>±1.12 | 27.45<br>±1.03 | 19.08<br>±1.11 |
| **7-14** | 26.44<br>±1.16 | 19.46<br>±1.83 | 27.19<br>±0.84 | 19.59<br>±1.29 | 27.84<sup>s1</sup><br>±2.12 | 19.39<br>±1.52 | 27.64<br>±0.84 | 19.00<br>±1.02 |
| **14-17** | 25.90<br>±0.89 | 19.27<br>±1.34 | 25.47<br>±1.83 | 19.40<br>±0.76 | 26.33<br>±2.71 | 18.47<br>±1.22 | 26.17<br>±0.82 | 18.73<br>±1.17 |
| **17-21** | 26.38<br>±0.97 | 19.88<br>±1.72 | 26.50<br>±0.77 | 20.08<br>±1.18 | 27.10<br>±2.31 | 19.35<br>±1.52 | 26.93<br>±0.86 | 19.13<br>±0.64 |
| **14-21** | 26.17<br>±0.93 | 19.61<br>±1.53 | 26.06<br>±1.19 | 19.79<br>±0.64 | 26.77<br>±2.42 | 18.97<br>±1.35 | 26.60<br>±0.67 | 18.96<br>±0.61 |
| **21-24** | 21.80<br>±0.77 | 15.90<br>±0.74 | 22.07<br>±1.03 | 16.23<br>±0.68 | 22.27<br>±1.61 | 15.97<br>±0.34 | 22.47<br>±0.66 | 15.63<br>±0.55 |
| **24-28** | 27.70<br>±1.04 | 20.70<br>±1.38 | 28.75<br>±1.21 | 21.33<br>±1.44 | 29.13<br>±1.92 | 21.08<br>±0.91 | 29.18<br>±1.28 | 20.45<br>±0.55 |
| **21-28** | 25.17<br>±0.89 | 18.64<br>±1.09 | 25.89<br>±1.10 | 19.14<br>±0.96 | 26.19<br>±1.73 | 18.89<br>±0.54 | 26.30<br>±0.83 | 18.39<br>±0.32 |
| **0-28** | 25.46<br>±0.91 | 18.89<br>±1.23 | 25.88<br>±0.87 | 19.05<br>±0.81 | 36.25<br>±1.58 | 18.79<br>±1.09 | 26.26<br>±0.90 | 18.44<br>±0.61 |

n=10 animals/sex/group
M=male
F=Femal
<sup>s1</sup> – p <0.05
<sup>s2</sup> – p <0.01
<sup>s3</sup> – p <0.001

**Table 6:** Summary of Mean Food Efficiency and Mean Daily Dietary Intake of LegH Prep – 28-Day Dietary Study.

| Mean Food Efficiency (mg/kg/day) |  |  |  |  |  |  |  |  | Mean Daily Dietary Intake of LegH Prep (mg/kg/day) |  |  |  |  |  |  |  |  |
| --- | --- | --- | --- | --- | --- | --- | --- | --- | --- | --- | --- | --- | --- | --- | --- | --- | --- |
| LegH Dose Levels | 0<br>(mg/kg/day) |  | 250<br>(mg/kg/day) |  | 500<br>(mg/kg/day) |  | 750<br>(mg/kg/day) |  | LegH Prep Dose Levels | 0<br>(mg/kg/day) |  | 512<br>(mg/kg/day) |  | 1024<br>(mg/kg/day) |  | 1536<br>(mg/kg/day) |  |
| Study Day | M | F | M | F | M | F | M | F | Study Days | M | F | M | F | M | F | M | F |
| **0-7** | 0.304<br>±0.046 | 0.193<br>±0.055 | 0.312<br>±0.038 | 0.215<br>±0.052 | 0.319<br>±0.051 | 0.226<br>±0.037 | 0.329<br>±0.046 | 0.206<br>±0.038 | **0-7** | 0.0<br>±0.0 | 0.0<br>±0.0 | 485.4<br>±20.9 | 498.0<br>±43.5 | 966.5<br>±42.5 | 995.9<br>±29.2 | 1459.8<br>±103.0 | 1481.1<br>±115.0 |
| **7-14** | 0.241<br>±0.025 | 0.148<br>±0.093 | 0.248<br>±0.043 | 0.128<br>±0.037 | 0.258<br>±0.041 | 0.146<br>±0.033 | 0.241<br>±0.040 | 0.165<br>±0.040 | **7-14** | 0.0<br>±0.0 | 0.0<br>±0.0 | 540.5<br>±24.5 | 541.9<br>±49.9 | 1095.9<br>±53.5 | 1064.6<br>±39.2 | 1631.5<br>±78.9 | 1604.6<br>±116.7 |
| **14-21** | 0.223<br>±0.044 | 0.149<br>±0.057 | 0.217<br>±0.022 | 0.077<sup>S2</sup><br>±0.042 | 0.227<br>±0.026 | 0.112<br>±0.049 | 0.217<br>±0.031 | 0.126<br>±0.052 | **14-21** | 0.0<br>±0.0 | 0.0<br>±0.0 | 503.2<br>±30.7 | 518.8<br>±53.9 | 1007.2<br>±61.7 | 1015.1<br>±34.3 | 1513.7<br>±81.1 | 1537.2<br>±92.3 |
| **21-28** | 0.121<br>±0.040 | 0.080<br>±0.052 | 0.123<br>±0.032 | 0.111<br>±0.041 | 0.139<br>±0.040 | 0.109<br>±0.033 | 0.139<br>±0.019 | 0.083<br>±0.044 | **21-28** | 0.0<br>±0.0 | 0.0<br>±0.0 | 495.9<br>±33.2 | 482.4<br>±41.9 | 973.0<br>±49.1 | 994.0<br>±56.0 | 1473.9<br>±92.5 | 1460.2<br>±79.0 |
| **0-28** | 0.222<br>±0.031 | 0.142<br>±0.027 | 0.224<br>±0.028 | 0.131<br>±0.028 | 0.235<br>±0.034 | 0.147<br>±0.011 | 0.230<br>±0.027 | 0.144<br>±0.019 | **0-28** | 0.0<br>±0.0 | 0.0<br>±0.0 | 478.9<br>±24.7 | 497.8<br>±42.8 | 954.7<br>±36.0 | 983.4<br>±29.0 | 1438.2<br>±78.6 | 1470.4<br>±88.2 |

n=10 animals/sex/group
M=male
F=female
<sup>S1</sup> – p <0.05
<sup>S2</sup> – p <0.01
<sup>S3</sup> – p <0.001

#### Pathology

There were to no test-substance-related changes in hematology parameters for male or rats (Table 7). Statically significant increase in red blood cell, hematocrit, and hemoglobin values and absolute basophil counts for Group 2 females and decreased absolute reticulocyte counts in Group 3 females were non-dose dependent and were interpreted to be within the expected biological variation and therefore not toxicologically relevant and not test article dependent (Table 7). There were no test-substance-related changes in coagulation parameters for female rats. A non-dose dependent increase in activated partial thromboplastin time (APPT) was observed in Group 3 and 4 males. Due to its very slight magnitude and lack of correlating pathological or clinical finding, this change is considered non-adverse. There were no test-substance-related changes in serum chemistry parameters for male rats (Table 8). Alkaline phosphatase (ALKP) was minimally decreased in a non-dose-dependent manner for Group 2 and Group 4 females (Table 8). This minimal decrease was not correlated with concurrent clinical pathology or histopathology changes and due to its limited clinical relevance is interpreted to have no toxicological significance and was not test-article dependent. Other differences in serum chemistry parameters that were statistically significant consisted of increased albumin and potassium values in Group 3 males, decreased glucose and chloride in Groups 2 and 3 females, increased globulin values in Group 3 females and increased calcium in Groups 2 and 3 females. These were generally of small magnitude, lacked a response in a dose-dependent manner and are interpreted to be within expected biological variation and considered to be of no toxicological relevance and non-test-article-dependent. There were no test-substance-related changes in urinalysis parameters for males or female rats. (Table 9)

**Table 7:** Hematology & Coagulation – 28-Day Dietary Study.

| Clinical Pathology - Hematology |  |  |  |  |  |  |  |  | Historical Means<br>And Ranges |  |
| --- | --- | --- | --- | --- | --- | --- | --- | --- | --- | --- |
| LegH Dose Levels | 0 (mg/kg/day) |  | 250 (mg/kg/day) |  | 500 (mg/kg/day) |  | 750 (mg/kg/day) |  |  |  |
| Parameter | M | F | M | F | M | F | M | F | M | F |
| RBC<br>(x106/mL) | 7.72<br>± 0.23 | 7.59<br>± 0.24 | 7.60<br>± 0.34 | 8.01<sup>S1</sup><br>± 0.38 | 7.61<br>± 0.35 | 7.86<br>± 0.24 | 7.70<br>± 0.27 | 7.63<br>± 0.30 | 8.75<br>5.07-10.04 | 8.26<br>6.99-9.34 |
| HGB<br>(g/dL) | 15.6<br>± 0.3 | 15.3<br>± 0.5 | 15.4<br>± 0.6 | 16.2<sup>S1</sup><br>± 0.5 | 15.5<br>± 0.6 | 15.7<br>± 0.4 | 15.9<br>± 0.4 | 15.5<br>± 0.6 | 15.7<br>10.5-17.4 | 15.4<br>13.4-17.3 |
| HCT<br>(%) | 45.5<br>± 0.9 | 43.6<br>± 1.2 | 45.1<br>± 1.5 | 45.9<sup>S1</sup><br>± 1.2 | 45.1<br>± 1.7 | 44.7<br>± 1.3 | 45.9<br>± 0.8 | 44.0<br>± 1.7 | 46.1<br>34.8-50.6 | 44.8<br>38.0-49.4 |
| MCV<br>(fL) | 58.9<br>± 1.0 | 57.5<br>± 1.1 | 59.3<br>± 2.3 | 57.4<br>± 2.2 | 59.3<br>± 1.5 | 56.8<br>± 1.2 | 59.7<br>± 1.9 | 57.7<br>± 2.2 | 52.8<br>47.5-68.6 | 54.3<br>49.9-58.6 |
| MCH<br>(pg) | 20.3<br>± 0.5 | 20.2<br>± 0.3 | 20.3<br>± 0.9 | 20.2<br>± 0.7 | 20.4<br>± 0.5 | 20.0<br>± 0.5 | 20.6<br>± 0.7 | 20.3<br>± 0.7 | 17.9<br>15.3-20.7 | 18.7<br>17.0-20.8 |
| MCHC<br>(g/dL) | 34.4<br>± 0.4 | 35.2<br>± 0.7 | 34.2<br>± 0.4 | 35.3<br>± 0.3 | 34.4<br>± 0.3 | 35.2<br>± 0.4 | 34.5<br>± 0.5 | 35.2<br>± 0.5 | 34.0<br>30.1-36.7 | 34.5<br>32.5-36.5 |
| RDW<br>(%) | 12.1<br>± 0.3 | 11.3<br>± 0.4 | 12.5<br>± 0.5 | 11.3<br>± 0.5 | 12.5<br>± 0.3 | 11.2<br>± 0.3 | 12.3<br>± 0.5 | 11.5<br>± 0.5 | 13.3<br>11.3-31.2 | 11.6<br>10.1-13.1 |
| PLT<br>(x103/mL) | 1160<br>± 121 | 1190<br>± 108 | 1202<br>± 69 | 1176<br>± 127 | 1171<br>± 76 | 1230<br>± 115 | 1227<br>± 185 | 1229<br>± 114 | 990<br>404-1799 | 1013<br>448-1594 |
| WBC<br>(x103/mL) | 13.00<br>± 1.33 | 10.08<br>± 1.70 | 14.41<br>± 2.67 | 11.87<br>± 1.75 | 11.13<br>± 1.82 | 11.59<br>± 3.35 | 13.45<br>± 4.41 | 10.19<br>± 3.72 | 11.94<br>2.75-22.23 | 7.66<br>2.41-17.04 |
| ANEU<br>(x103/mL) | 1.91<br>± 0.67 | 1.48<br>± 0.30 | 1.99<br>± 0.43 | 1.56<br>± 0.58 | 1.75<br>± 0.43 | 1.68<br>± 0.85 | 1.57<br>± 0.62 | 1.54<br>± 1.10 | 2.11<br>0.53-9.39 | 1.08<br>0.31-4.08 |
| ALYM<br>(x103/mL) | 10.49<br>± 1.17 | 8.15<br>± 1.58 | 11.79<br>± 2.48 | 9.74<br>± 1.43 | 8.86<br>± 1.70 | 9.29<br>± 2.71 | 11.29<br>± 4.15 | 8.21<br>± 2.88 | 9.12<br>1.47-20.32 | 6.16<br>1.79-12.84 |
| AMON<br>(x103/mL) | 0.31<br>± 0.10 | 0.25<br>± 0.15 | 0.34<br>± 0.11 | 0.29<br>± 0.06 | 0.28<br>± 0.05 | 0.33<br>± 0.15 | 0.30<br>± 0.10 | 0.22<br>± 0.14 | 0.35<br>0.09-0.93 | 0.20<br>0.03-0.56 |
| AEOS<br>(x103/mL) | 0.12<br>± 0.04 | 0.11<br>± 0.03 | 0.13<br>± 0.08 | 0.13<br>± 0.04 | 0.11<br>± 0.04 | 0.15<br>± 0.05 | 0.11<br>± 0.05 | 0.12<br>± 0.06 | 0.17<br>0.00-0.88 | 0.13<br>0.04-0.84 |
| ABAS<br>(x103/mL) | 0.09<br>± 0.03 | 0.04<br>± 0.01 | 0.09<br>± 0.04 | 0.07<sup>S1</sup><br>± 0.03 | 0.07<br>± 0.02 | 0.06<br>± 0.03 | 0.10<br>± 0.06 | 0.05<br>± 0.04 | 0.06<br>0.00-0.27 | 0.03<br>0.00-0.15 |
| ALUC<br>(x103/mL) | 0.08<br>± 0.03 | 0.05<br>± 0.02 | 0.08<br>± 0.03 | 0.07<br>± 0.02 | 0.06<br>± 0.02 | 0.07<br>± 0.03 | 0.08<br>± 0.04 | 0.05<br>± 0.04 | 0.11<br>0.00-0.47 | 0.06<br>0.00-0.26 |
| ARET<br>(x103/mL) | 232.6<br>± 31.2 | 205.8<br>± 33.9 | 235.8<br>± 40.7 | 182.4<br>± 32.9 | 246.3<br>± 24.1 | 169.1<sup>S1</sup><br>± 30.9 | 243.8<br>± 41.1 | 184.2<br>± 33.7 | 219.5<br>98.6-1913.0 | 164.6<br>27.7-277.2 |
| PT<br>(sec) | 10.7<br>± 0.3 | 10.0<br>± 0.2 | 10.7<br>± 0.4 | 9.8<br>± 0.2 | 10.6<br>± 0.2 | 10.0<br>± 0.3 | 10.6<br>± 0.2 | 9.8<br>± 0.2 | 10.53<br>9.5-12.1 | 9.9<br>9.2-10.7 |
| APTT<br>(sec) | 20.2<br>± 2.4 | 21.9<br>± 2.5 | 23.8<br>± 5.3 | 20.0<br>± 3.1 | 24.9<sup>S1</sup><br>± 6.9 | 20.8<br>± 5.0 | 24.9<sup>S1</sup><br>± 6.9 | 19.4<br>± 1.9 | 19.86<br>13.5-54.8 | 19.0<br>13.2-48.5 |

n=10 animals/sex/group
M=male
F=female
<sup>S1</sup> – p <0.05
<sup>S2</sup> – p <0.01
<sup>S3</sup> – p <0.001

**Table 8:**
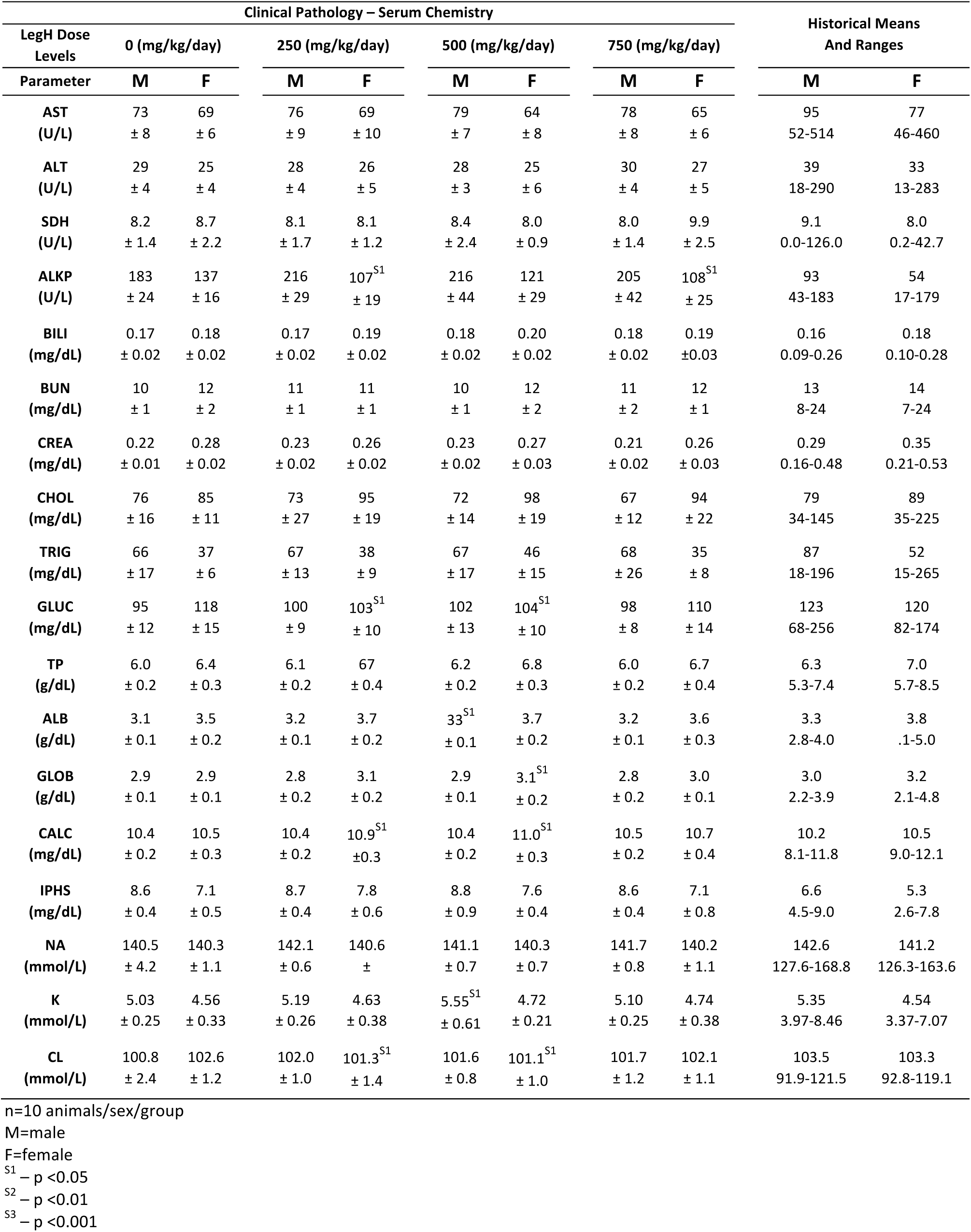
Serum Chemistry28-Day Dietary Study.

**Table 9:** Urinalysis – 28-Day Dietary Study.

| Clinical Pathology – Urinalysis |  |  |  |  |  |  |  |  |  | Historical Means<br>And Ranges |
| --- | --- | --- | --- | --- | --- | --- | --- | --- | --- | --- |
| LegH Dose<br>Levels | 0 (mg/kg/day) |  | 250 (mg/kg/day) |  | 500 (mg/kg/day) |  | 750 (mg/kg/day) |  |  |  |
| Parameter | M | F | M | F | M | F | M | F | M | F |
| UVOL<br>(mL) | 11.7<br>± 8.2 | 7.8<br>± 6.4 | 11.5<br>± 9.8 | 6.8<br>± 5.1 | 12.3<br>± 7.3 | 6.5<br>± 3.0 | 14.3<br>± 7.7 | 6.6<br>± 4.1 | 7.3<br>0.3-36.0 | 5.4<br>0.1-39.0 |
| pH | 6.5<br>± 0.3 | 6.4<br>± 0.4 | 6.5<br>± 0.4 | 6.2<br>± 0.4 | 6.6<br>± 0.4 | 6.6<br>± 0.6 | 6.6<br>± 0.4 | 6.5<br>± 0.6 | 6.4<br>5.0-8.5 | 6.5<br>5.0-8.5 |
| SG | 1.027<br>± 0.019 | 1.037<br>± 0.027 | 1.027<br>± 0.015 | 1.035<br>± 0.023 | 1.026<br>± 0.015 | 1.028<br>± 0.011 | 1.024<br>± 0.019 | 1.030<br>± 0.013 | 1.052<br>1.007-1.100 | 1.047<br>1.009-1.100 |
| URO<br>(EU/dL) | 0.03<br>± 0.03 | 0.2<br>± 0.0 | 0.2<br>± 0.0 | 0.2<br>± 0.0 | 0.3<br>± 0.3 | 0.2<br>± 0.0 | 0.2<br>± 0.0 | 0.3<br>± 0.3 | 0.3<br>0.2-1.0 | 0.3<br>0.2-1.0 |
| UMTP<br>(mg/dL) | 104<br>± 49 | 43<br>± 34 | 241<br>± 265 | 41<br>± 25 | 124<br>± 80 | 34<br>± 12 | 111<br>± 97 | 44<br>± 30 | 213<br>18-1330 | 92<br>6-7400 |

n=10 animals/sex/group
M=male
F=female
<sup>s1</sup> – p <0.05
<sup>s2</sup> – p <0.01
<sup>s3</sup> – p <0.001

There were no test-article-dependent effects observed during necropsy, organ weights, macroscopic evaluation and microscopic evaluation in male and female rats, with a single exception of a distinct estrous cycle stage distribution in the female rats. The estrous cycle consists of four stages: proestrus, estrus, metestrus, and diestrous. Each stage has characteristic reproductive organ weights and pathology. At study termination, Group 2 and 4 females had an increased incidence of metestrus and a decreased incidence of estrus compared to Groups 1 and 3. Consistent with the estrous cycle stage distribution, Group 2 and 4 females also had decreased presence of fluid-filled uteri and dilated uterine lumens and decreased uterine weights compared to Group 1 and 3 females (Table 10-11). These decreases did not correlate with adverse histopathological findings and are therefore interpreted to be non-adverse. The presence of both new and old ovarian corpora lutea in females from all groups indicated that all females were cycling normally. All other microscopic findings at the study day 29/30 time point were also unrelated to administration of LegH Prep and can be observed in the age and strain of rats used in this study.^63,64^ Although the differences in estrous cycle stage distribution between groups was likely due sampling and assessing estrous cycle distribution on a single day, rather than using a longitudinal study, a more extensive and rigorous longitudinal study was performed focusing on the potential effect of LegH Prep on the estrous cycle.

**Table 10:**
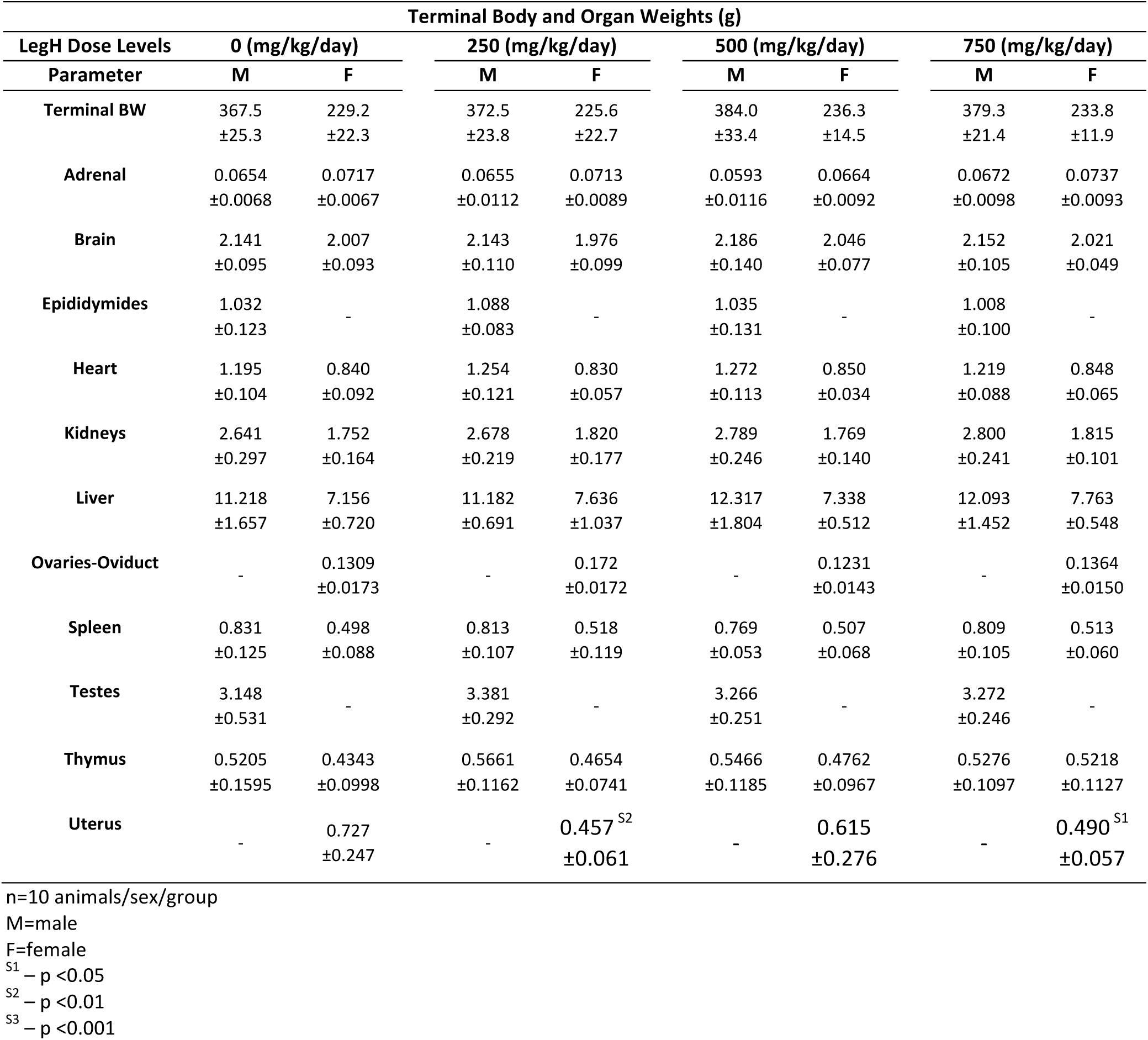
Summary of Mean Terminal Body Weights and Organ Weights – 28-Day Dietary Study.

| Terminal Body and Organ Weights (g) |  |  |  |  |  |  |  |  |
| --- | --- | --- | --- | --- | --- | --- | --- | --- |
| LegH Dose Levels | 0 (mg/kg/day) |  | 250 (mg/kg/day) |  | 500 (mg/kg/day) |  | 750 (mg/kg/day) |  |
| Parameter | M | F | M | F | M | F | M | F |
| Terminal BW | 367.5 | 229.2 | 372.5 | 225.6 | 384.0 | 236.3 | 379.3 | 233.8 |
|  | ±25.3 | ±22.3 | ±23.8 | ±22.7 | ±33.4 | ±14.5 | ±21.4 | ±11.9 |
| Adrenal | 0.0654 | 0.0717 | 0.0655 | 0.0713 | 0.0593 | 0.0664 | 0.0672 | 0.0737 |
|  | ±0.0068 | ±0.0067 | ±0.0112 | ±0.0089 | ±0.0116 | ±0.0092 | ±0.0098 | ±0.0093 |
| Brain | 2.141 | 2.007 | 2.143 | 1.976 | 2.186 | 2.046 | 2.152 | 2.021 |
|  | ±0.095 | ±0.093 | ±0.110 | ±0.099 | ±0.140 | ±0.077 | ±0.105 | ±0.049 |
| Epididymides | 1.032 | - | 1.088 | - | 1.035 | - | 1.008 | - |
|  | ±0.123 |  | ±0.083 |  | ±0.131 |  | ±0.100 |  |
| Heart | 1.195 | 0.840 | 1.254 | 0.830 | 1.272 | 0.850 | 1.219 | 0.848 |
|  | ±0.104 | ±0.092 | ±0.121 | ±0.057 | ±0.113 | ±0.034 | ±0.088 | ±0.065 |
| Kidneys | 2.641 | 1.752 | 2.678 | 1.820 | 2.789 | 1.769 | 2.800 | 1.815 |
|  | ±0.297 | ±0.164 | ±0.219 | ±0.177 | ±0.246 | ±0.140 | ±0.241 | ±0.101 |
| Liver | 11.218 | 7.156 | 11.182 | 7.636 | 12.317 | 7.338 | 12.093 | 7.763 |
|  | ±1.657 | ±0.720 | ±0.691 | ±1.037 | ±1.804 | ±0.512 | ±1.452 | ±0.548 |
| Ovaries-Oviduct | - | 0.1309 | - | 0.172 | - | 0.1231 | - | 0.1364 |
|  |  | ±0.0173 |  | ±0.0172 |  | ±0.0143 |  | ±0.0150 |
| Spleen | 0.831 | 0.498 | 0.813 | 0.518 | 0.769 | 0.507 | 0.809 | 0.513 |
|  | ±0.125 | ±0.088 | ±0.107 | ±0.119 | ±0.053 | ±0.068 | ±0.105 | ±0.060 |
| Testes | 3.148 | - | 3.381 | - | 3.266 | - | 3.272 | - |
|  | ±0.531 |  | ±0.292 |  | ±0.251 |  | ±0.246 |  |
| Thymus | 0.5205 | 0.4343 | 0.5661 | 0.4654 | 0.5466 | 0.4762 | 0.5276 | 0.5218 |
|  | ±0.1595 | ±0.0998 | ±0.1162 | ±0.0741 | ±0.1185 | ±0.0967 | ±0.1097 | ±0.1127 |
| Uterus | - | 0.727 | - | 0.457<sup>s2</sup> | - | 0.615 | - | 0.490<sup>s1</sup> |
|  |  | ±0.247 |  | ±0.061 |  | ±0.276 |  | ±0.057 |

n=10 animals/sex/group
M=male
F=female
<sup>S1</sup> – p <0.05
<sup>S2</sup> – p <0.01
<sup>S3</sup> – p <0.001

**Table 11:** Summary of Mean Relative Organ-to-Body Weights – 28-Day Dietary Study.

| Organ-to-Body Weight Ratios (g) |  |  |  |  |  |  |  |  |
| --- | --- | --- | --- | --- | --- | --- | --- | --- |
| LegH Dose Levels | 0 (mg/kg/day) |  | 250 (mg/kg/day) |  | 500 (mg/kg/day) |  | 750 (mg/kg/day) |  |
| Parameter | M | F | M | F | M | F | M | F |
| **Adrenal / TBW** | 0.1781<br>±0.0165 | 0.3139<br>±0.0265 | 0.1766<br>±0.0328 | 0.3168<br>±0.0336 | 0.1540<br>±0.0253 | 0.2812<br>±0.0372 | 0.1773<br>±0.0264 | 0.3157<br>±0.0399 |
| **Brain / TBW** | 5.846<br>±0.411 | 8.801<br>±0.545 | 5.766<br>±0.355 | 8.828<br>±0.852 | 5.722<br>±0.497 | 8.692<br>±0.686 | 5.682<br>±0.294 | 8.664<br>±0.492 |
| **Epididymides / TBW** | 2.8075<br>±0.2682 | - | 2.9351<br>±0.3125 | - | 2.7030<br>±0.3143 | - | 2.6712<br>±0.3544 | - |
| **Heart / TBW** | 3.251<br>±0.151 | 3.665<br>±0.189 | 3.362<br>±0.151 | 3.692<br>±0.171 | 3.315<br>±0.128 | 3.605<br>±0.178 | 3.214<br>±0.149 | 3.625<br>±0.163 |
| **Kidneys / TBW** | 7.184<br>±0.610 | 7.657<br>±0.412 | 7.199<br>±0.541 | 8.094<br>±0.639 | 7.274<br>±0.421 | 7.505<br>±0.657 | 7.387<br>±0.560 | 7.783<br>±0.602 |
| **Liver / TBW** | 30.549<br>±4.348 | 31.278<br>±2.212 | 30.052<br>±1.405 | 33.819<br>±2.693 | 31.962<br>±2.654 | 31.158<br>±2.883 | 31.839<br>±3.559 | 33.269<br>±2.772 |
| **Ovaries-Oviduct / TBW** | - | 0.5727<br>±0.0669 | - | 0.5635<br>±0.0474 | - | 0.5222<br>±0.0643 | - | 0.5835<br>±0.0581 |
| **Spleen / TBW** | 2.256<br>±0.255 | 2.171<br>±0.300 | 2.199<br>±0.391 | 2.284<br>±0.384 | 2.012<br>±0.184 | 2.149<br>±0.291 | 2.139<br>±0.312 | 2.191<br>±0.206 |
| **Testes / TBW** | 8.549<br>±1.201 | - | 9.108<br>±0.971 | - | 8.564<br>±0.970 | - | 8.657<br>±0.885 | - |
| **Thymus / TBW** | 1.4134<br>±0.4037 | 1.8863<br>±0.3463 | 1.5209<br>±0.3105 | 2.0742<br>±0.3287 | 1.4171<br>±0.2319 | 2.0184<br>±0.4057 | 1.3939<br>±0.2919 | 2.2362<br>±0.4918 |
| **Uterus / TBW** | - | 3.159<br>±0.949 | - | 2.060<sup>S2</sup><br>±0.452 | - | 2.579<br>±1.063 | - | 2.103<sup>S2</sup><br>±0.277 |

n=10 animals/sex/group
M=male
F=female
<sup>S1</sup> – p <0.05
<sup>S2</sup> – p <0.01
<sup>S3</sup> – p <0.001

### 28-Day Investigative Study in Rats with a 14-Day Pre-Dosing Estrous Cycle Determination

A 28-day dietary feeding study was performed with female rats to thoroughly evaluate the estrous cycle stage distributions observed in the previous 28-day dietary feeding study. To ensure all animals had normal estrous cyclicity prior to the 28-day dosing phase, estrous cycle stage was determined daily for all animals for 14 days. Additionally, estrous cycle stage was determined for all animals for the last 14 days of the 28-day dosing period. At study termination, reproductive organs were analyzed. Administered doses of 0, 512, 1024 and 1536 mg/kg/day of freeze-dried LegH Prep correspond to 0, 250, 500 and 750 mg/kg/day of LegH, respectively. The mean overall daily intake of the test substance in Group 2, 3 and 4 female rats was 250, 496, and 738 mg/kg/day LegH, respectively. The animals are considered to have received acceptable dose levels.

#### Mortality, clinical signs, body weight/food consumption

No mortalities were observed during this study. There were no clinical observations attributable to the administration of LegH Prep. There were no body weight, body weight gain, food consumption, or food efficiency findings considered attributable to LegH Prep administration with the exception of a single incidental increase (p<0.05) in mean daily body weight, mean food consumption and mean food efficiency for Group 2 animals on days 21-28. This increase was transient, non-dose-dependent, and interpreted to have no toxicological relevance.

#### Estrous cycle evaluation and Pathology

There were no test-substance-related changes in average estrus length attributable to LegH Prep administration (Table 12). There were no macroscopic or microscopic findings related to the administration of LegH Prep. A single Group 2 animal had prolonged estrus based on morphology of the ovaries (large atretic follicles, multiple corpus lutea at a similar state of atresia) and presence of squamous metaplasia of the uterus. These findings were considered spontaneous and incidental due to the lack of similar findings at higher dose levels. One Group 1 animal had large atretic follicles observed in both ovaries, and one Group 4 animal had lutenized follicles (follicles with evidence of lutenization in the wall but which have not ovulated) in both ovaries. Both of these observations are reported as background findings in rats of the strain and age used in this study^65^ and were considered incidental because of their singular occurrences. There were no test-substance-related changes in absolute or relative reproductive organ weight values in female rats treated with LegH Prep (Table 13). Longitudinal daily monitoring of estrous cycle stage demonstrated that, despite intrinsically normal estrous cycles, the distribution of estrous cycle stages on any given day can be markedly different from the within-rat distribution over time (Figure 1).

**Table 12:**
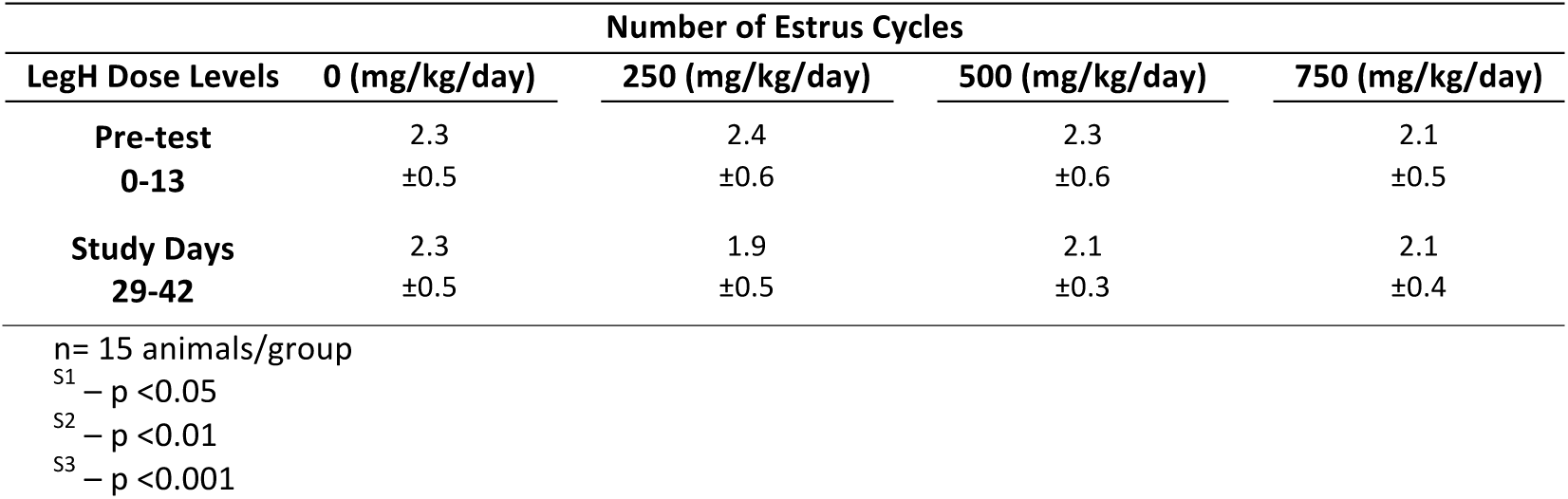
Estrus Cycles – 28-Day Dietary Study with Pre-Dosing Estrous Cycle Determination.

**Table 13:**
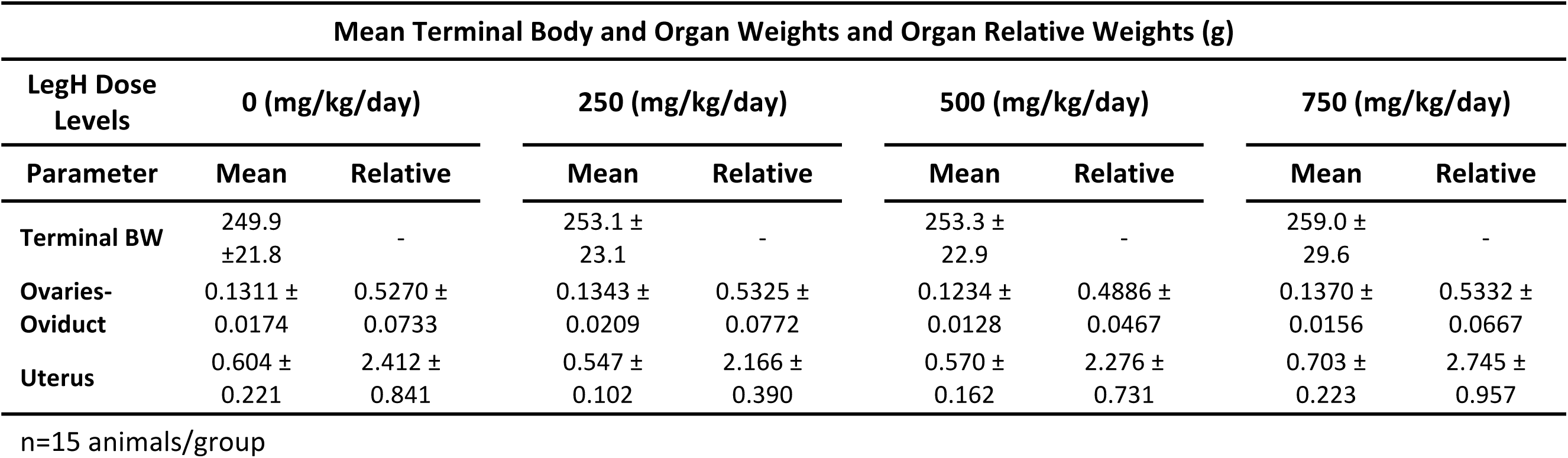
Summary of Mean Terminal Body and Organ Weights and Organ Relative Weights– 28-Day Dietary Study with Pre-Dosing Estrous Cycle Determination.

**Figure 1.**
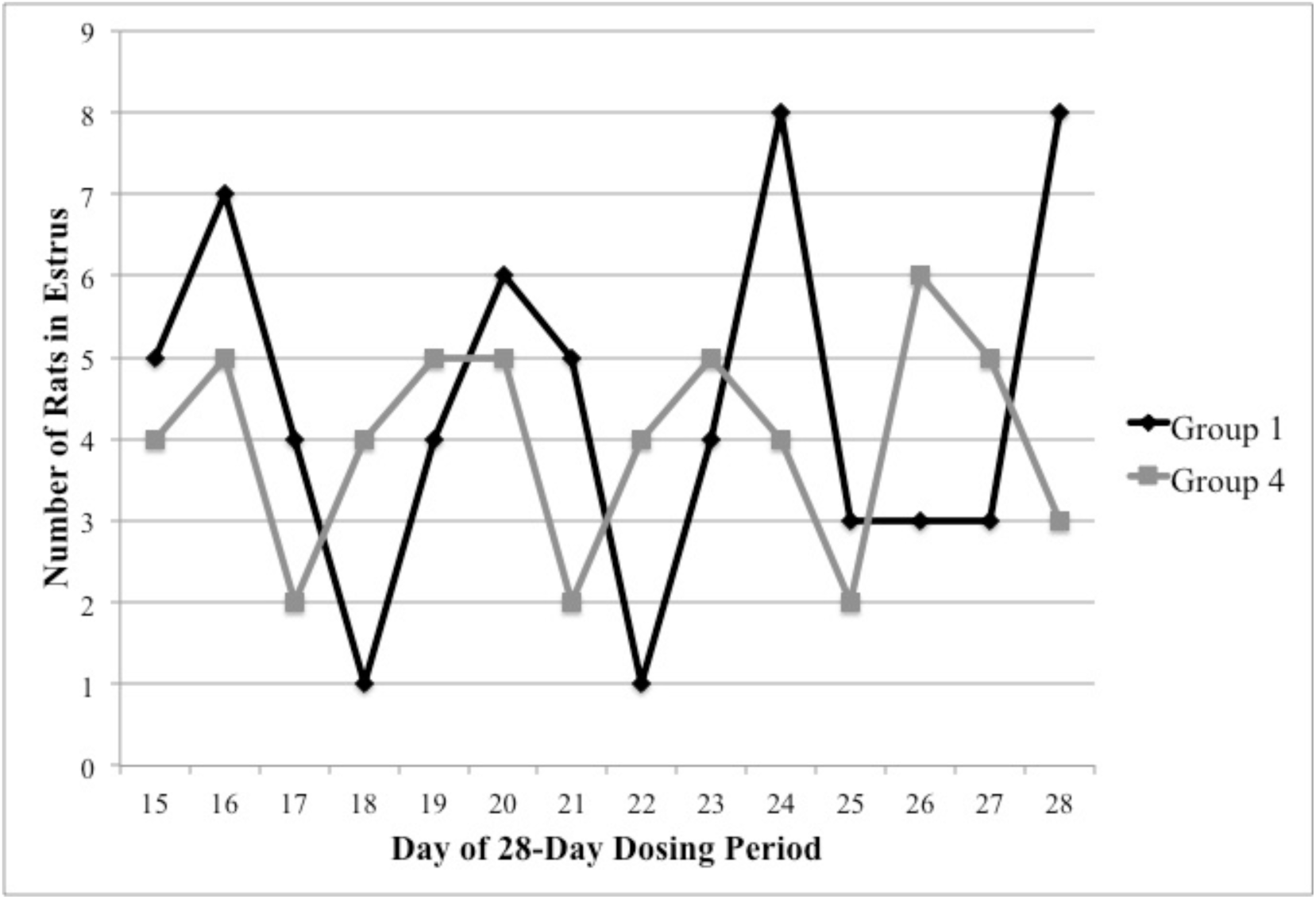
Number of rats in the estrus phase of the estrous cycle for the first 10 rats within Groups 1 and 4 on each day. Data are from the 28-day dietary study with pre-dosing estrous cycle determination.

### Pathology Peer Review

Because there were no test-article-dependent effects observed in the estrous cycle study, a pathology peer review was performed on the initial 28-day dietary feeding study. The review pathologist evaluated histopathology in all female reproductive organs and corresponding macroscopic and microscopic observations noted by the study pathologist. Following the peer review, the study pathologist and review pathologist reached a consensus that there were no test-article-dependent effects on the female estrous cycle and reproductive organs.

## Discussion

Heme is ubiquitous in the human diet and has been consumed for thousands of years. Replacing the myoglobin that catalyzes the unique flavor chemistry of meat derived from animals with LegH from soy opens an opportunity to develop plant-based meats that deliver to consumers the pleasure they demand from animal-derived meats, with a small fraction of the environmental impact. To evaluate the safety profile of soy leghemoglobin produced in Pichia, we conducted a series of *in vitro* and *in vivo* studies.

The Pichia-derived LegH Prep was non-mutagenic in the bacterial reverse mutation test, which evaluated five strains of bacteria and eight different concentrations of LegH Prep up to a maximum dose of 5000 μg LegH/plate. Similarly, LegH Prep was non-clastogenic in the chromosomal aberration test, which evaluated chromosomal rearrangements in HPBL following 4 hr (with and without metabolic activation) and 24 hr (without metabolic activation) incubations with LegH Prep. These assays tested LegH concentrations up to 5000 μg/ml for the 4 hr incubations. Due to test article precipitation and decreased percent mitotic index, 1000 μg/ml LegH was the maximum dose evaluated for the 24 hr incubation. Together, these results demonstrate that LegH Prep is non-mutagenic and non-clastogenic under the *in vitro* conditions tested.

To evaluate the *in vivo* safety profile for potential systemic toxicity, a 28-day feeding study was conducted in rats in which LegH Prep was administered in the diet. Animals were monitored for clinical observations, food consumption, body weight, ophthalmology, clinical pathology, necropsy and histopathology. There were no LegH Prep-dependent effects observed with the exception of a distinct estrous cycle stage distribution in Group 2 and 4 females at study termination. Group 2 and 4 females had an increased incidence of the metestrus stage of the estrous cycle, decreased presence of fluid filled uteri and dilated uterine lumens, and decreased uterine weight compared to Group 1 and 3 females (Table 10-11). However, the correlation between estrous cycle stage and reproductive organ weight and pathology for each animal was consistent with published literature on normal healthy rats.^66^ Therefore, although the estrous cycle stage distribution was different between groups, there were no data to suggest an adverse impact on the health of the female animals; the presence of both new and old ovarian corpora lutea indicated normal estrous cyclicity.^65^ Without evidence of an adverse effect in the female ovary or uterus pathology, the decrease in relative and absolute uterine weights in Group 2 and 4 females was interpreted to be non-adverse. Moreover, decreased uterine weight is normal for animals in the metestrus stage of the estrous cycle.^67^

An in-depth follow-up 28-day dietary feeding study in female rats was performed with longitudinal estrous cycle monitoring and evaluation of reproductive organ weights, gross necropsy, and histopathology. The results demonstrated that LegH Prep had no impact on the estrous cycle length, distribution, or female reproductive organ health (Table 12-13). Despite intrinsically normal estrous cycles, different estrous cycle stage distributions were observed between groups on any given day (Figure 1). This highlights the importance of longitudinal estrous cycle monitoring to evaluate estrous cyclicity. For example, if the estrous cycle stages were only monitored on a single day, a completely different conclusion would have been drawn regarding the test article-effect on estrous cycle if the animals had been analyzed, for example, on day 18 of the dosing period compared to day 21 (Figure 1). The single day sampling artifact readily accounts for the increased incidence of metestrus observed in the initial 28-day dietary feeding study. Moreover, a pathology peer review of the original 28-day study resulted in a consensus between the study pathologist and review pathologist that LegH Prep did not affect the estrous cycle.

Together, these systemic toxicity and reproductive health feeding studies in rats established a NOAEL of 750 mg/kg/day LegH for both sexes, which was the maximum dose administered. Collectively these *in vitro* and *in vivo* results establish that LegH Prep, containing both soy leghemoglobin protein and Pichia proteins from the production host, is safe for its intended use in ground beef analogue products.

Creating safe, delicious plant-based meats to replace animal-derived meats in the diet is critical to reducing and eventually eliminating the environmental impact of the animal farming industry. Impossible Foods Inc. has shown that plant-based meat containing up to 0.8% LegH delivers flavors and aromas that are characteristic of animal-derived meat.^11^ This study established a NOAEL of 750 mg/kg/day LegH, which is over 100 times higher than the 90^th^ percentile EDI. This maximum dose is equivalent to an average sized person (60 kg) consuming 5625g (12 lbs) of plant-based ground beef analogue with 0.8% LegH per day. Thus, LegH Prep is safe for its intended use in ground beef analogue products.

## Authorship

RZF, SK, MS and OM designed the experiments; RZF, MS, and PA performed the experiments; RZF, MS, PA, SK, and OM analyzed the data; RZF, MS, and OM wrote the manuscript.

## Acknowledgements

We acknowledge ELT and KM for assistance with initial drafting of the manuscript; POB, ELT, GLY, and JLV for critical evaluation of the manuscript; MP and NH for feedback on study design; CT for critical review of chromosome aberration test; and HY and PA for analytical chemistry support.

## Footnotes

a Datamonitor estimates the US meat analogue volume was 53M kg in 2009

